# Sexual antagonism leads to a mosaic of X–autosome conflict

**DOI:** 10.1101/746032

**Authors:** Steven A. Frank, Manus M. Patten

## Abstract

Males and females have different optimal values for some traits, such as body size. When the same genes control these traits in both sexes, selection pushes in opposite directions in males and females. Alleles at autosomal loci spend equal amounts of time in males and females, suggesting that the sexually antagonistic selective forces may approximately balance between the opposing optima. Frank and Crespi noted that alleles on the X chromosome spend twice as much time in diploid females as in haploid males. That distinction between the sexes may tend to favor X-linked genes that push more strongly toward the female optimum than the male optimum. The female bias of X-linked genes conflicts with the intermediate optimum of autosomal genes, potentially creating an X-autosome conflict. Patten has recently argued that explicit genetic assumptions about dominance and the relative magnitude of allelic effects may lead X-linked genes to favor the male rather than the female optimum, contradicting Frank and Crespi. This article combines the insights of those prior analyses into a new, more general theory. We find some parameter combinations for X-linked loci that favor a female bias and other parameter combinations that favor a male bias. We conclude that the X likely contains a mosaic pattern of loci that conflict with autosomes over sexually antagonistic traits. The overall tendency for a female or male bias on the X depends on prior assumptions about the distribution of key parameters across X-linked loci. Those parameters include the dominance coefficient and the way in which ploidy influences the magnitude of allelic effects.

## Introduction

Sexual antagonism arises when males and females have different optimum values for a trait with a shared genetic basis. Selection in males pushes the evolution of the trait in one direction, and selection in females pushes in the other direction (Rice & Chippindale, 2001; Bonduriansky & Chenoweth, 2009; van Doorn, 2009). For autosomal alleles, which occur equally in males and females, the opposing male and female selective pressures tend to balance, and we expect intermediate phenotypes.

How do opposing selective forces in males and females play out on the X chromosome? Females are diploid on the X, with two alleles at each locus. Males are haploid, with one allele at each locus. Frank & Crespi (2011) argued that selection will tend to push more strongly toward the female optimum, because each allele spends two-thirds of its time in females and one-third of its time in males. The resulting tendency toward female-biased optima on the X conflicts with the tendency toward intermediate phenotypes favored by autosomes.

Patten (2019) argued that selection on the X may favor traits that tend toward the male optimum rather than the female optimum. His conclusion follows from two factors. First, dominance can mask some mutations on diploid female X chromosomes but not on haploid male X chromosomes. Second, if the effect per locus on trait expression is the same in females as in males, then each allele in a diploid female has half the effect of an allele in a haploid male.

Each of these assumptions reduces the genetic variance in fitness in females, weakening the overall selective pressure that females impose on trait evolution relative to males. In consequence, trait evolution may tend toward the male optimum.

These contrasting conclusions about the direction of sex bias favored by X-linked genes arise from different assumptions about the genetics of traits. In this article, we develop a more general theory that subsumes the prior models. From that more general analysis, we show why X-linked loci may vary with regard to the direction of sex bias. Thus, the X may be a mosaic of loci that conflict in different ways with the intermediate tendency for trait evolution caused by autosomal loci.

Mosaic sex bias along the X is interesting because X-autosome conflict potentially plays a role in disease (Frank & Crespi, 2011), in speciation (Crespi & Nosil, 2013), and in genomic evolution (Patten, 2018). With regard to the specific genetic assumptions, X-autosome conflict depends in interesting ways on the variability among loci in dominance and the relation between ploidy level and the contribution of each allele to phenotypic value.

We first develop a simple model that highlights how the key genetic parameters influence the bias of the X-linked genes toward the male or female optimum. We then consider how the distribution of genetic parameter values determines the mosaicism in bias of the X and whether the overall bias is toward the male or the female optimum.

## Analysis

Frank & Crespi (2011) noted that two genetic factors can influence the dynamics of selection on the X chromosome under sexual antagonism. First, females are diploid on the X, whereas males are haploid. Thus, X-linked alleles spend twice as much time in females as in males, which can lead to a greater potential for selection in females relative to males. For example, when females have a double dose of the same allele in homozygotes, that potentially increases the effect of the locus on trait values relative to the same hemizygous locus in males. Second, alleles in females at loci that are heterozygous may be hidden from expression if recessive, weakening the relative selection pressure in females (Vicoso & Charlesworth, 2006). In males, all alleles are potentially exposed to selection because they are not paired with another allele at the same locus.

Frank & Crespi (2011) did not analyze the role of dominant versus recessive allelic effects in females. Instead, they argued that most traits of interest would be influenced by many genetic loci. The alleles at each of those many loci would tend to have a small and more or less additive effect on the overall trait value. In addition, epistatic interactions between loci may contribute more strongly to any non-additivity than dominance within loci, such that haploid males and diploid females do not experience greatly differing nonadditivity and masking of rare alleles. As shown in the following analysis, these assumptions favor a female bias on the X.

Patten (2019) extended the theory by explicitly including a dominance parameter. In addition, Patten (2019), following Rice (1984), assumed that each female homozygous locus and male hemizygous locus have equivalent phenotypic consequences. By contrast, Frank & Crespi (2011) assumed that, at a subset of loci, a female homozygote has a greater phenotypic effect than a male hemizygote. Here, we analyze a more general model that subsumes the earlier work and clarifies the different assumptions in the prior analyses.

### Parameters

Suppose an X-linked locus is fixed for the *X*_1_ allele, with *X*_1_*X*_1_ females and *X*_1_*Y* males. Consider the fitness of a sexually antagonistic mutant *X*_2_ allele that increases fitness in one sex and decreases fitness in the other. Let mutant males, *X*_2_*Y*, have fitness 1 *+ M*. Homozygous mutant females, *X*_2_*X*_2_, have fitness 1 − *γM*, with *γ >* 0. If *M* is positive, the mutant has beneficial effects in males and deleterious effects in females. The opposite effects hold for negative *M*. Heterozygous mutant females, *X*_1_*X*_2_, have fitness 1 − *hγM*, in which 0 *≤ h ≤* 1 is the standard dominance coefficient.

The parameter *γ = αδ* has two separate components. First, *α >* 0 scales fitness effects in females relative to males, such that the mutant effect is *M* in males and −*αM* in females. Second, 1 *≤ δ ≤* 2, scales the effect of a diploid female locus relative to a haploid male locus.

Rice (1984) and Patten (2019) assumed equal per locus effects in females and males, *δ =* 1, which means that each allele in a female has, on average, one-half the effect of each allele in a male, or, equivalently, that the lone allele in a male has twice the effect of each allele in a female. Frank & Crespi (2011) assumed equal per allele effects in females and males, *δ =* 2, which means that each allele has the same effect in females and males. Here, we generalize the analysis by specifying the continuous parameter, *δ*, between those two endpoints.

### Invasion of a rare mutant

For *M >* 0, a rare male-beneficial mutant invades when (Parsons, 1961)

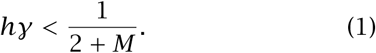

When this condition holds for a sufficiently large fraction of new mutations, then X-linked loci will be biased toward the male optimum.

For *M <* 0, a rare female-beneficial mutant allele, *X*_2_, invades and spreads in a population fixed for *X*_1_, when

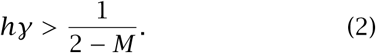

When this condition holds for a sufficiently large fraction of new mutations, then X-linked loci will be biased toward the female optimum.

### Sex bias of individual X-linked loci

If new mutations at a locus are biased toward the optimum favored by one sex, then that locus will tend to push in the direction of the favored sex. Mutational bias can arise at both autosomal and X-linked loci.

Several additional factors uniquely influence sex bias at X-linked loci. First, if new mutants at a locus are biased with regard to dominance, *h*, then that bias can lead to the favoring of one sex over the other. Dominant mutations, *h >* 1*/*2, favor a female bias, and recessive mutations, *h <* 1*/*2, favor a male bias. The greater the deviation of *h* from 1*/*2, the greater the expected bias.

Second, larger fitness effects of mutants, with greater absolute *M* values, favor a male bias. That bias arises because the condition for male beneficial mutants in eqn 1 becomes relatively easier to satisfy for increasing *M* than does the condition for female beneficial mutants in eqn 2 for decreasing values of *M*.

Third, in the composite parameter *γ = αδ*, the parameter *δ* is the female:male ratio for the per locus contribution to fitness. An increase in *δ* enhances the relative selective intensity acting on females relative to males, with the consequence of increasing the tendency of X-linked loci to favor the female optimum.

Simplifying assumptions provide further insight about the conditions for sex bias. Suppose that mutations have symmetric effects on females and males, *α =* 1, and that mutations have relatively small effects on fitness, *M* → 0. Then the condition for a male-beneficial mutation (*M >* 0) to increase is

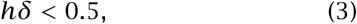

and the condition for a female-beneficial mutation (*M* < 0) to increase is

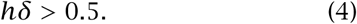

If we assume that allelic effects are additive, that is, neither dominant nor recessive, with *h =* 0.5, then the condition for male bias, *δ <* 1 is never satisfied. The condition for female bias is *δ >* 1. If *δ =* 1, then there is no bias.

The parameter *δ* takes on its minimal value of one when a mutant allele in a female has one-half the effect of a mutant allele in a male. If the effect in a female is greater than one-half that in males, then *δ >* 1. When mutant alleles have the same additive effect in females and males, then *δ =* 2. Assuming that at least some loci have *δ >* 1, the bias is always toward females for loci that experience new mutations with small additive effects (Frank & Crespi, 2011).

By contrast, when *δ =* 1, a male bias can arise under particular assumptions about dominance. Suppose, for example, that fitness is a bell-shaped function of phenotype in both sexes, such that mutations benefitting females (*M* < 0) are dominant and mutations harming females (*M* > 0) are recessive because of the curvature of the fitness surface (Connallon & Chenoweth, 2019). Patten (2019) showed that such reversals of dominance can lead to a male bias for sexually antagonistic traits at X-linked loci.

Finally, dosage compensation by X inactivation causes females to be effectively haploid. With males and females effectively haploid, there is no sex-associated asymmetry on the X, and therefore no inherent tendency for the X to favor one sex over the other.

However, as noted by Frank & Crespi (2011), “About 15% of genes on the human X chromosome escape inactivation, and another 10% of X-linked loci are variably expressed on inactive X chromosomes (Carrel & Willard, 2005). Thus, a significant number of X-linked loci may be expressed from both copies and may conflict with autosomes. Occasional diploid expression on the X is sufficient to create the conflict.” Under systems with X inactivation, our theory applies to the many loci that at least partially escape inactivation.

## Conclusion

Variation in parameter values across loci means that, inevitably, some loci will have a male bias and other loci will have a female bias. The X chromosome will therefore be mosaic for the direction of sex biases and X-autosome conflict, with some X-linked loci pushing toward the female optimum and other loci pushing toward the male optimum.

Across the entire X chromosome, the overall tendency for a bias toward females or males depends on the distribution of parameter values. If mutations tend to have small additive effects (*M* → 0 and *h* ≈ 1*/*2), as may happen for the sort of polygenic traits likely to be influenced by sexual antagonism, then the overall bias when *δ >* 1 is strongly toward females (Frank & Crespi, 2011). By contrast, if new mutations tend be recessive *(h <* 1*/*2), have sufficiently large fitness effects (*M* » 0), or have an association between dominance and fitness effects, then the overall bias when *δ =* 1 is likely to be toward males (Patten, 2019).

In summary, a simple genetic model and broadly reasonable assumptions lead to mosaicism in X-autosome conflict over sexually antagonistic traits.

## Acknowledgments

The Donald Bren Foundation supports SAF’s research. SAF completed this work while on sabbatical in the Theoretical Biology group of the Institute for Integrative Biology at ETH Zürich.

## Author contributions

SAF initiated this research. SAF and MMP jointly worked out the differences in their prior models, formulated the final form of the analysis, and wrote the manuscript.

